# Differentiating shame- and guilt-proneness with heart rate variability in Chinese young adults

**DOI:** 10.1101/2023.03.02.530780

**Authors:** Isaac N. Ip, Fiona N. Y. Ching, Hey Tou Chiu, Ariel H. Y. Keung, Savio W. H. Wong

## Abstract

A high proneness to experience shame and guilt has been associated with psychopathology. Despite their similarity, shame- and guilt-proneness have different psychological and neurobiological correlates. The present study aims to compare the physiological correlates between shame- and guilt-proneness. Resting heart rate variability (HRV), a peripheral biomarker of emotion dysregulation and psychopathology, was measured in a sample of 60 Chinese young adults with two sessions of electrocardiogram recording. Proneness to shame and guilt were measured by the Test of Self-Conscious Affect 3. Hierarchical linear modeling indicated that guilt-proneness was positively associated with HRV while shame-proneness was not. Our findings implied that shame- and guilt-proneness have different relations with HRV. The distinct physiological relations are discussed with respect to the adaptive/maladaptive nature of shame- and guilt-proneness.

---

Both shame and guilt are fundamental moral, self-conscious emotions that serve to motivate us to act in accordance with social norms and personal standards (Else-Quest et al., 2012). When we are caught lying to a loved one, for example, a sense of shame or guilt helps us appraise the situation negatively. Shame, on the one hand, involves a painful sense of shrinking and worthlessness i.e. a global devaluation of the self (Tangney & Dearing, 2004), so that we would be more likely to avoid committing the same mistake. Guilt, on the other hand, is characterized by a sense of tension and remorse that involves a preoccupation with the transgression or failure (Tangney & Dearing, 2004), which would make us feel compelled to rectify the situation. Despite their important functions in regulating social behaviors, a high proneness to experience shame and guilt across various situations can be detrimental to wellbeing and associated with psychopathology, especially in adolescence and young adulthood (Muris & Meesters, 2014). Meta-analytic studies have shown that predisposition to experience shame and guilt were strongly associated with anxiety disorders (Cândea & Szentagotai-Tătar, 2018) and depression (Kim et al., 2011).

Resting heart rate variability (HRV), a peripheral measure of autonomic regulation, has emerged as a non-invasive biomarker of emotion dysregulation and psychopathology (Beauchaine, 2015). Meta-analyses indicated that lower levels of HRV are associated with anxiety and depression. For instance, across various types of anxiety disorders, including panic disorder, post-traumatic stress disorder, generalized anxiety disorder, and social anxiety disorder, patients displayed reductions in HRV relative to controls (Chalmers et al., 2014). Patients with depression also showed lower HRV than healthy controls, and higher levels of depression severity was associated with lower HRV (Kemp et al., 2010). Conversely, higher levels of resting HRV is associated with more flexible emotional responding and adaptive emotion regulation in healthy adults (see Balzarotti et al., 2017, for a review). This link between autonomic and emotion regulation can be explained at the levels of the peripheral and central nervous systems. At the peripheral level, a robust parasympathetic/vagal regulation of heart rate, which is directly supported by the functioning of the myelinated vagus, can quickly mobilize or calm an individual’s physiological arousal and is crucial for adaptive emotion regulation (Porges, 2007). At the central level, processes of cognitive, emotional, and physiological regulation are supported by a common neural circuit that involves the inhibitory circuits of the prefrontal cortex (Thayer & Lane, 2009). Lower HRV may therefore indicate poor functioning of the vagus nerve or disruptions to the prefrontal inhibitory circuits, resulting in difficulties in emotion regulation.

The observed relations between psychopathology and proneness to shame and guilt on the one hand and HRV on the other suggest a possible link between proneness to shame and guilt and HRV. Feelings of shame and guilt involve self-awareness and self-representations i.e. these emotions arise only when individuals become aware that they have failed to live up to some actual or ideal self-representations (Tracy & Robins, 2004). These self-referential processes, which underlie self-conscious emotions such as shame and guilt, are supported by the medial prefrontal cortex (mPFC) (Sebastian et al., 2008). Neuroimaging studies that contrasted activations during self-conscious emotions to basic emotions or a neutral condition consistently found that the mPFC was involved in processing self-conscious emotions (e.g. Burnett et al., 2009; Gilead et al., 2016; Takahashi et al., 2004). Apart from self-referential processes, the mPFC has been associated with parasympathetic regulation. Wong et al. (2007) showed that deactivation in the ventral portion of mPFC was associated with vagal withdrawal, indicated by an increase in heart rate, during a graded handgrip exercise; stronger deactivation correlated with a larger increase in heart rate. These neuroimaging findings provide a neurophysiological basis for the possible link between proneness to shame and guilt and HRV.

Although shame and guilt appear to be very similar, researchers have argued that there are meaningful differences between the two emotions on phenomenological, cognitive, and motivational levels. In terms of cognitive appraisal, shame is generated by stable, uncontrollable, and global attributions (“I’m not good enough”), whereas guilt is generated by unstable, controllable, and specific attributions (“I did not try hard enough”) of an eliciting event (Tracy & Robins, 2004). Consistent with this distinction, research has shown that people who were more prone to shame tended to attribute failures to their own character or incapability (aspects of the self that is global, stable and uncontrollable), whereas people who were more prone to guilt tended to attribute failures to lack of effort or specific behaviors that are unstable and controllable (Tracy & Robins, 2006). In terms of motivation, shame is often associated with action tendency to withdrawing from the situation, while guilt is associated with motivation to repair (Cohen et al., 2011). These differences have led researchers to suggest that, although shame and guilt are both negatively valenced, guilt has relatively more adaptive facets than shame (Tangney & Dearing, 2004). Findings from meta-analyses support this differentiation. For instance, two subtypes of guilt-proneness, namely generalized guilt and contextual-maladaptive guilt, were as strongly associated with anxiety and depression as shame-proneness was (Cândea & Szentagotai-Tătar, 2018; Kim et al., 2011). However, the overall association of anxiety/depression with guilt-proneness was weaker than that with shame-proneness. Evidently, this is because a more adaptive subtype of guilt-proneness, contextual-legitimate guilt, only had a weak association with psychopathology. This subtype of guilt-proneness is characterized by accurate and realistic attribution of responsibility and thus is regarded as situationally appropriate.

Differences between shame- and guilt-proneness have also been reported in neuroimaging studies. On the one hand, higher levels of proneness to shame, measured by the Experience of Shame Scale, was associated with smaller amygdala and thinner posterior cingulate cortex (PCC) in healthy adolescents and young adults (Whittle et al., 2016). On the other hand, children who were prone to pathological guilt had smaller anterior insula (Belden et al., 2015), while larger anterior cingulate cortex (ACC) was associated with compensatory behavior in a task designed to induce guilt in participants (Yu et al., 2014). These neuroimaging findings show that proneness to shame and guilt not only have different psychological properties, but also neuroanatomical correlates despite their apparent similarity. Interestingly, although emotion theories and research typically have a strong emphasis on the involvement of physiological states, research on shame and guilt to date have focused solely on biological correlates at the cortical level, thus leaving peripheral physiology a research area largely unexplored.

We therefore aimed to investigate whether proneness to shame and guilt would be associated with individual differences in HRV. First, based on the observation that a high proneness to experience shame and guilt and lower levels of HRV are both associated with psychopathology, we hypothesized that shame- and guilt-proneness would be negatively associated with HRV. In addition, given that proneness to shame and guilt can be differentiated by their psychological and neural correlates, we would like to explore whether this hypothesized association with HRV would differ between shame- and guilt-proneness. Specifically, in view of the meta-analytic findings that the association of anxiety/depression with guilt-proneness was weaker than that with shame-proneness, we speculated that the extent of negative association of HRV with guilt-proneness would also be less than that with shame-proneness.

## Methods

### Participants and Procedures

Participants were 60 Chinese young adults (28 females; age: *M* = 21.6, *SD* = 3.10) recruited by convenience sampling through an advertisement posted on the university mass email. Based on an effect size distribution analysis of 297 HRV effect sizes (Quintana, 2017), 61 participants are needed to achieve 80% power to detect a medium effect size. Exclusion criteria were smokers, known blood pressure conditions, and chronic heart issues or respiratory conditions. Inclusion criterion was 18-35 years of age. One participant was taking medications that potentially influence the autonomic nervous system (ANS) and therefore was excluded from data analyses. The rest of them were not on any medication. All participants reported no psychological illness.

Before their visit, participants were reminded by email to refrain from food and caffeine for two hours and alcohol and intensive physical activity for 24 hours prior to the study. At the beginning of the study, the participants gave informed consent and confirmed whether they had abided by the instructions. They were then cabled by a same-sex experimenter and were seated in a comfortable chair in front of a workstation. After completing the first part of a questionnaire during a habituation period, they underwent a HRV measurement for six minutes. The participants were instructed to remain seated, relax and breathe spontaneously while looking at a fixation on the screen during the measurement. They then completed the second part of the questionnaire, followed by another HRV measurement with the same procedures as the first one. The study received ethical approval from the local institutional review board.

### Measures

#### Shame- and Guilt-Proneness

Proneness to shame and guilt were measured by the Test of Self-Conscious Affect 3 (TOSCA-3; Tangney & Dearing, 2004). The TOSCA-3 is a scenario-based measure of self-conscious emotions consisting of 16 scenarios and response sets (e.g. “You are driving down the road and you hit a small animal: 1. You would think: ‘I’m terrible.’ (*Shame*); 2. You’d feel bad you hadn’t been more alert driving down the road.” (*Guilt*)). Responses are rated on a scale from 1 (“not likely”) to 5 (“very likely”). A Chinese version of TOSCA-3 was used (Gao et al., 2013).^1^ Subscales of shame- and guilt-proneness both demonstrated good reliability (*α*s: shame = .791, guilt = .756).

#### Social Desirability

As responses in TOSCA-3 can be susceptible to social desirability in the Asian population (Hasui et al., 2009), we also measured social desirability using the Marlowe-Crowne Social Desirability Scale short form C (MCSDS-C; Reynolds, 1982). The MCSDS-C is a 13-item scale that assesses how susceptible the participants’ responses are to social approval. Participants indicated whether each of the items (e.g. “I have never intensely disliked anyone.”) is true or false for themselves. A Chinese version of the scale was used (Liao, 2002; *α* = .543).

### Physiological Data Acquisition and Preprocessing

As reliability of standard short-term HRV measurements can be low and potentially pose a problem when HRV is assessed as a trait (Uhlig et al., 2020), we deliberately collected two measurements from each participant to account for situational specific variance (Bertsch et al., 2012; Laborde et al., 2017). Electrocardiogram (ECG) signals were sampled at 1000 Hz, transduced and amplified through a Biopac MP150 Bioamplifier and recorded by AcqKnowledge (Biopac Systems, USA). The ECG recordings were obtained using the Lead-II configuration with three standard Ag/AgCl electrodes (two below the left and right clavicle, one on the left lower ribcage).

Preprocessing of ECG data was based on a 5-minute interval between the first and last 30 seconds of the recordings. First, artifacts were detected by visual examination of the ECG time series. R-peak false-positive or false-negative detections were manually corrected; ectopic beats were identified. Next, interbeat intervals (IBI) from the ECG were extracted. The artifacts were then removed, and values were estimated via cubic spline interpolation of neighboring IBIs. Finally, frequency domain measures of HRV were calculated using ARTiiFACT (Kaufmann et al., 2011). The absolute and relative power of the high frequency band (i.e. HFabs and HFn.u., respectively), which reflect vagal influences on heart rate (Laborde et al., 2017), were used for data analyses. As expected, the two indices were highly correlated with each other in the first and second measurement (*r* = .725, *p* < .001, and *r* = .560, *p* < .001, respectively). Both indices demonstrated good reliability in our sample, having strong correlations across the two measurements (HFabs: *r* = .928, *p* < .001; HFn.u.: *r* = .717, *p* < .001).

### Data Analyses

Descriptive statistics of background variables, social desirability, shame- and guilt-proneness, and HRV indices were first calculated. HFabs was expectedly highly skewed and was log-transformed before bivariate correlations were calculated. Given that HRV data were nested within participants, hierarchical linear modeling (HLM) was used to test our hypotheses. Specifically, we used random intercepts models to test the associations between HRV and proneness to shame and guilt. Given that HRV is influenced by a number of physiological factors including BMI, it is necessary to include BMI into the model as a covariate (Laborde et al., 2017). Gender was also included as a covariate because of its correlations with HRV (Koenig & Thayer, 2016). Social desirability was not included in the model due to its low reliability and lack of correlations with shame- or guilt-proneness in our sample. Following Tangney and Dearing’s (2004) methodology, shame- and guilt-proneness were entered into the same model to test their unique associations with HRV.

## Results

### Bivariate Correlations

Bivariate correlations and descriptive statistics are shown in Table 1. Among background variables, female participants (coded as 0) were significantly higher than male participants (coded as 1) in HFn.u. of the first and second measurements (*r* = -.399, *p* = .002, and *r* = -.408, *p* = .001, respectively). Age, BMI, and social desirability were not correlated with outcome variables.

**Table 1.**
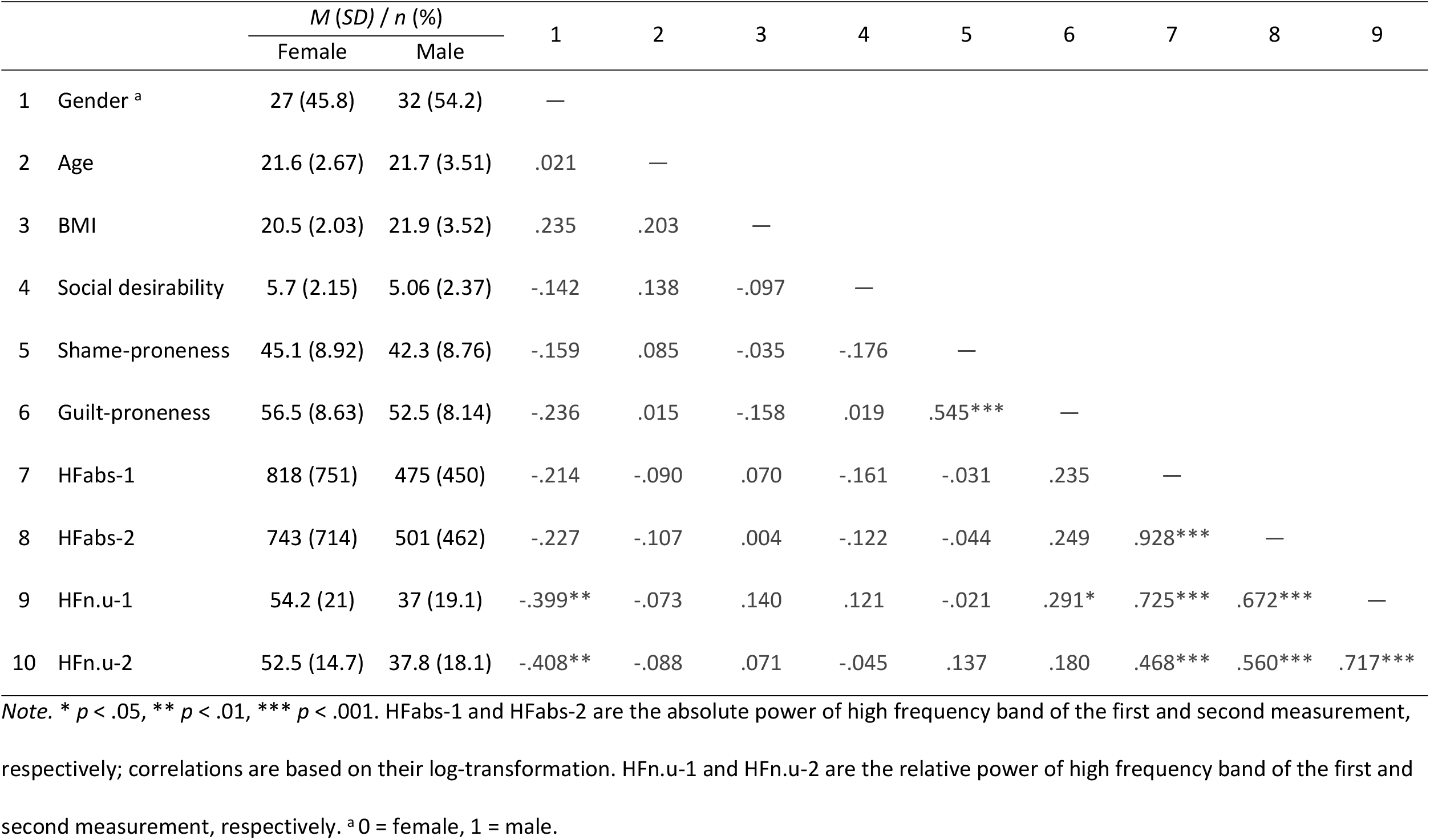
Descriptive Statistics and Bivariate Correlations for Background and Outcome Variables

As expected, shame-proneness was positively related to guilt-proneness (*r* = .545, *p* < .001), replicating previous findings based on the TOSCA-3 (Gao et al., 2013; Tangney & Dearing, 2004). HFn.u. of the first measurement was positively correlated with guilt-proneness (*r* = .291, *p* = .026), while none of the HRV indices of the first or second measurement were correlated with shame-proneness.

### HLM

Results of the HLM are shown in Table 2. Compared to the null model, the overall fit of the model consisting of BMI, gender, shame- and guilt-proneness was significantly improved in predicting lnHFabs (Δχ^2^ (4 d.f.) = 9.74, *p* < .05) and HFn.u. (Δχ^2^ (4 d.f.) = 20.43, *p* <. 01). Controlling for BMI, gender, and shame-proneness, guilt-proneness was significantly positively associated with lnHFabs (*B* = 0.05, *S*.*E*. = 0.02, *t* = 2.46, *p* = .017) and HFn.u. (*B* = 0.61, *S*.*E*. = 0.29, *t* = 2.10, *p* = .040). Meanwhile, controlling for BMI, gender, and guilt-proneness, shame-proneness was not associated with lnHFabs (*B* = -0.03, *S*.*E*. = 0.02, *t* = - 1.84, *p* = .071) and HFn.u. (*B* = -0.34, *S*.*E*. = 0.27, *t* = -1.23, *p* = .223).

**Table 2.**
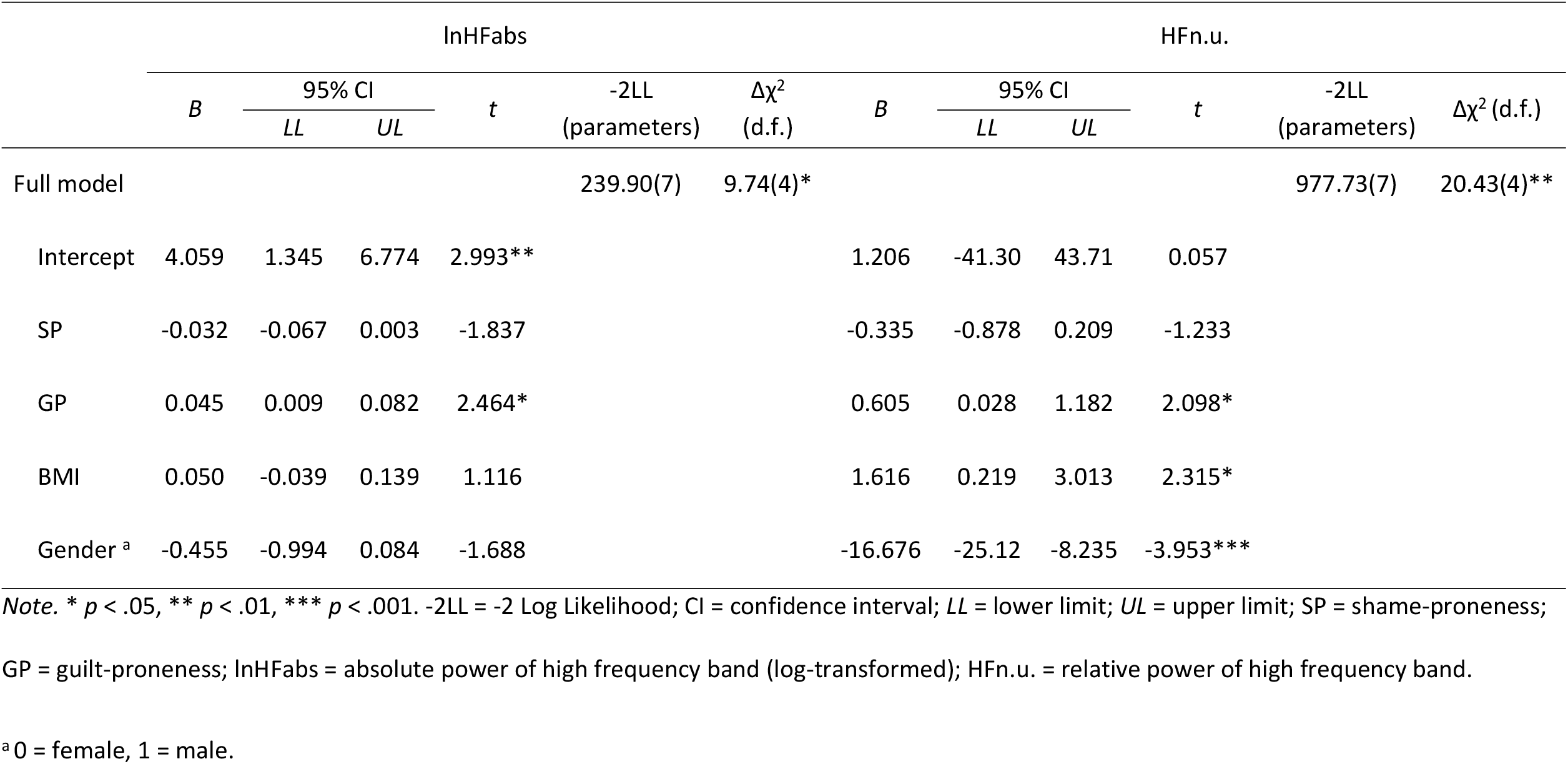
Multilevel Random Intercept Models Predicting HRV with Shame- and Guilt-Proneness, Controlling for BMI and Gender

## Discussion

The current study investigated whether shame- and guilt-proneness would be differentially associated with resting HRV in a group of healthy Chinese young adults. Our results indicated that neither shame- nor guilt-proneness were negatively associated with HRV. Instead, guilt-proneness was positively associated with HRV, whereas shame-proneness was largely unassociated with HRV. This implies that shame- and guilt-proneness have different relations with HRV in Chinese young adults, although the direction of their associations was contrary to our hypotheses.

The finding that shame- and guilt-proneness have different relations with HRV aligns with the argument that meaningful differences exist between shame and guilt. As mentioned, previous psychological and neuroimaging research support this distinction by showing that shame- and guilt-proneness can be differentiated by their cognitive appraisals and neurobiological correlates. Our findings provide further evidence at the level of peripheral physiology for this distinction.

At first glance, the finding that guilt-proneness was positively associated with HRV seems to be counterintuitive, given the empirical evidence available showing that proneness to guilt is positively associated with psychopathology. A possible explanation for this finding may pertain to the fact that some facets of guilt-proneness are more adaptive than others. As discussed, meta-analyses showed that the more adaptive subtype of guilt-proneness, contextual-legitimate guilt, only had a weak association with psychopathology, which contributed to the overall weaker association of anxiety/depression with guilt-proneness than that with shame-proneness. This situationally-appropriate subtype of guilt-proneness relies upon adaptive emotion regulation so that individuals can make accurate and realistic attribution of responsibility rationally when confronted by their own transgression or failure. In that case, a positive association would be observed between this particular subtype of guilt-proneness and HRV.

The present study assessed guilt- and shame-proneness using the TOSCA-3, which is arguably the most widely-used instrument for measuring shame- and guilt-proneness concurrently. Nonetheless, it has been suggested that the guilt-proneness items in TOSCA-3 tend to measure adaptive social behaviors such as reparative behaviors that can be motivated by guilt. Luyten et al., (2002) analyzed the items of TOSCA and found that maladaptive aspects of guilt were underrepresented. For instance, rumination and anxiety associated with guilt were only measured by one item each, while generalized guilt was not assessed. Thus, we may have observed this finding because the instrument that we used tended to measure more adaptive aspects of guilt-proneness rather than its maladaptive facets.

Nevertheless, it is unclear why a negative association between shame-proneness and HRV was not observed in our data. We speculate that the relationship between HRV and shame-proneness may vary across cultural contexts. For instance, Wong and Tsai (2007) argued that shame can be experienced differently in cultural contexts that promote an independent versus an interdependent self. Specifically, shame would be associated with maladaptive consequences in the former context and more adaptive consequences in the latter. Hence, while we did not find a clear relationship between shame-proneness and HRV in a cultural context that presumably promote an interdependent self, it is possible that studies conducted in another cultural context (e.g. one that promote an independent self) may yield different results. Future cross-cultural studies should be able to test this hypothesis.

It should be noted that although HRV was measured twice for each participant, our study essentially used a cross-sectional design so causal relations cannot be warranted. Nonetheless, our findings suggest that shame- and guilt-proneness have different relations with HRV, an emerging biomarker of emotion dysregulation. To the best of our knowledge, this is the first study that investigated the physiological correlates of shame- and guilt-proneness. Our findings potentially contribute to the literature that a physiological correlate, namely HRV, may differentiate shame- and guilt-proneness and are consistent with the position that there are tangible differences between shame and guilt. Future studies are encouraged to elucidate their relations by including measures of proneness to shame and guilt developed from other theoretical frameworks as well as participants from more diverse cultural contexts.

## Footnotes

1 Since it had been translated for validation using samples from northern China, minor adjustments to the wordings were made to aid the participants in Hong Kong in comprehension.

